# Genome-wide analysis tracks the emergence of intraspecific polyploids in *Phragmites australis*

**DOI:** 10.1101/2021.09.05.458733

**Authors:** Cui Wang, Lele Liu, Meiqi Yin, Franziska Eller, Hans Brix, Tong Wang, Jarkko Salojärvi, Weihua Guo

**Author notes:** **Correspondence to:** Professor Weihua Guo, Assistant Professor Jarkko Salojärvi.

## Abstract

Polyploidization is a common event in plant evolution, and it plays an important role in plant speciation and adaptation. To address the role of polyploidization in grass diversification, we studied *Phragmites australis*, a species with intraspecific variation of chromosome numbers ranging from 2n=36 to 144. A combined analysis of genome structure, phylogeny and population genetics were used to study the evolution of *P. australis*. Whole-genome sequencing of three representative lineages revealed the allopolyploid origin of the species, with subgenome divergence dating back to approximately 29 million years ago, and the genomes showed hallmarks of relaxed selection associated with asexual propagation. Genome-wide analysis of 88 individuals from different populations around the world using restriction site associated DNA sequencing (RAD-seq) identified seven main intraspecific lineages with extensive genetic admixture. Each lineage was characterized by a distinct ploidy level, mostly tetraploid or octoploid, suggesting several polyploid events. Furthermore, we observed octoploid and hexaploid lineages at contact zones in Romania, Hungary and South Africa, suggestively due to genomic conflicts in allotetraploid parental lineages. Polyploidy may have evolved as a strategy to escape from the evolutionary dead-end of asexual propagation and the resulting decrease in genomic plasticity.

## Introduction

The rapidly increasing number of high quality plant genome assemblies have uncovered numerous polyploidization or whole genome duplication (WGD) events immediately prior to or co-occurring with the time of species divergence (Alix, et al. 2017). These events, originating from autopolyploid or allopolyploid hybrids, are particularly common among grasses (Estep, et al. 2014), for example, all cereals have undergone rho, sigma and tau WGD events. The grass family (Poaceae), which includes more than 10,000 species today, originated from a common ancestor with five chromosomes (Salse, et al. 2008). In cereals, a series of genome duplication and consecutive genome fractionation events, involving chromosomal structural rearrangements and losses of redundancy, led to a common ancestral genome with 12 chromosomes, shared across the early diverging grass subfamilies Anomochlooideae, Pharoideae, and Puelioideae (Levy and Feldman 2002; Salse, et al. 2008). Hence, all extant Poaceae species are paleopolyploids, in which polyploidization occurred millions of years ago (Levy and Feldman 2002). Depending on their lineages, the current diploids have experienced further WGDs in the form of allo- or autopolyploidization, giving rise to the current grass species.

A recent polyploidization event can be directly inferred when closely related species show different chromosome counts or genome sizes, which differ by an integer-valued factor. This can often be observed in species within the same genus, but in some cases different ploidy levels have been observed also within the same species, such as *Dupontia fisheri, Betula pendula, Lygeum spartum, Cynodon dactylon* and *Phragmites australis* (Raicu, et al. 1972; Brysting, et al. 2004; Salojärvi, et al. 2017; Abdeddaim-Boughanmi, et al. 2019; Zhang, et al. 2020). These species are thus ideal models to understand the factors that drive polyploidization. One critical step to characterize the extent of polyploidization is to assess the ploidy level directly by counting chromosome numbers of individual metaphases, or quantifying the genome size in the germplasm using, for example, flow cytometry and comparing it to the size in a known diploid representative. However, the tedious procedures and the need of appropriate facilities can be challenging. With the advance of next generation sequencing (NGS) technology, large quantities of NGS reads now provide a chance to infer the ploidy levels through alleles of the genetic markers. Tools such as ploidyNGS (dos Santos, et al. 2016), ConPADE (Margarido and Heckerman 2015) and nQuire (Weiß, et al. 2018) have been proposed for inferring ploidy levels by fitting a model to the allele frequencies from mapping against the reference genome or calculating the frequency of haplotypic contigs. However, the methods are still under development and demonstrate relatively high error rates. Therefore a robust method to determine the ploidy level is still needed to understand the evolution of polyploidy in plants.

Common reed, *Phragmites australis* (Cav.) Trin. ex Steud., is a tenacious cosmopolitan wetland species which provides habitats for small animals and is considered an ecosystem engineer. Similar to many other grass species, *Phragmites* has a basal chromosome count of x=12. However, the euploid chromosome count has proved to be versatile, ranging from 3x to 12x (3x, 4x, 6x, 7x, 8x, 10x, 11x and 12x), or even mixoploid (Connor, et al. 1998; Clevering and Lissner 1999). Tetraploid and octoploid individuals are most common in nature (Connor, et al. 1998; Clevering and Lissner 1999). Substantial efforts to investigate the evolution of speciation in *Phragmites* have been made, but the evolution of ploidy levels within this species is still unclear (Saltonstall 2002; Lambertini, et al. 2006; Lambertini, Mendelssohn, et al. 2012; Tanaka, et al. 2017; Liu, et al. 2020). To date, seven species, *P. australis, P. mauritianus* Kunth, *P. frutescens* H. Scholz, *P. dioica* Hackel ex Conert, *P. berlandieri* E. Fourn., *P. japonicus* Steudel and *P. vallatoria* (Plunk. ex L.) Veldk, have been described in *Phragmites* (Lambertini, et al. 2006). Most of them are distributed regionally, with only *P. australis* being cosmopolitan (Greuter and Scholz 1996; Clevering and Lissner 1999; Lambertini, et al. 2006). Ploidy variation was found in several species, e.g., *P. vallatoria* has 2x, 3x, 4x with aneuploid representatives at all three ploidy levels, *P. japonicus* preserves 4x and aneuploid 8x, and *P. mauritianus* is mostly 4x (Lambertini, et al. 2006). Large intraspecific variation was documented in *P. australis* by morphological traits and molecular markers, and at least three subspecies have been suggested: *P. australis ssp. australis, P. australis ssp. altissimus*, and *P. australis ssp. Americanus* (Peterson and Soreng 2004; Saltonstall, et al. 2004; Lambertini, et al. 2006).

Based on non-coding chloroplast regions *trnT-trnL* and *rbcL-psaI*, researchers have classified lineages to 57 haplotypes (Saltonstall 2016), roughly corresponding to geographic regions of origin. According to the standard naming system (Saltonstall 2002), several phylogenetic studies have found haplotype M to be commonly distributed in North America, Europe, and Asia, and later confirmed it to be an invasive lineage in North America (Saltonstall 2002). Haplotype I is distributed across South America, southern Pacific islands and Asia; haplotypes U and Q occur in Asia and Australia, whereas haplotype O is mainly distributed in northern China and haplotype P in Korea and middle to southern China. A number of unique haplotypes have emerged endemically in Mexico, or North America, and in southwestern China it is possible to find three main haplotypes I, Q, U (Saltonstall 2002; An, et al. 2012; Colin and Eguiarte 2016; Tanaka, et al. 2017). Because the chloroplast haplotypes are not continuously distributed within the same geographic region, it is difficult to conclude how they spread originally, especially when considering the variation observed in intraspecific ploidy levels. Studies based on amplified fragment length polymorphisms (AFLP) and microsatellites have provided further support to the geographic lineage classifications. However, the AFLP produced only 107 variable sites among the species, which may not be sufficient for analyzing allopolyploid species with up to 144 chromosomes (Lambertini, et al. 2006).

In United States, the range of exotic *Phragmites australis* has expanded from a very limited area to span almost the entire country within just 50 years, forcing the native lineages to the very north; it is thus considered an invasive plant (Saltonstall 2002). One hypothesis for the development of invasiveness in *P. australis* is related to polyploidizations (Clevering and Lissner 1999; Lambertini 2019), because polyploids, due to their higher level of genomic plasticity, can outperform their diploid progenitors in environments with high abiotic stress, and thus have access to a wider range of habitats (Godfree, et al. 2017). Successful allopolyploids may thus develop into invasive species and drive diploid ancestors to extinction by recurrent hybridizations with them.

In this study, we investigated the intraspecific genetic divergence, population genetic structure, historical demographics, and hybridization of *P. australis* using RAD-seq approaches that incorporated data from the nuclear genome and chloroplast from 88 individuals sampled throughout the geographic range of the species. To understand the geographic distribution of ploidy variation in the species, we developed a method based on alternative allele frequency of reads mapped to the reference genome to predict the ploidy level of each individual. We obtained high coverage Illumina sequencing data for the whole genomes of three individuals representing three lineages and assembled their draft genomes to investigate the genome evolution associated with polyploidization, and to give insights into the rise of whole genome duplications in grass species.

## Results and discussion

### Genome assemblies confirm allopolyploid structure of P. australis

We carried out RAD-sequencing for a collection of 88 *P. australis* individuals from Eurasian, North American, Oceanian and African continents (Fig. 1, Table 1). A subset of these samples was quantified with flow cytometry to obtain genome size estimate (Table 1; flow cytometry info provided by H. B., the samples have been used in previous studies (Lambertini, et al. 2006)). For initial exploration of the RAD-seq data we carried out a *de novo* analysis using Stacks (Catchen, et al. 2013). Even though the method assumes the organisms to be diploid and only considers biallelic sites, we were able to obtain enough qualified variant sites to describe the population grouping of the species.

**Figure 1.**
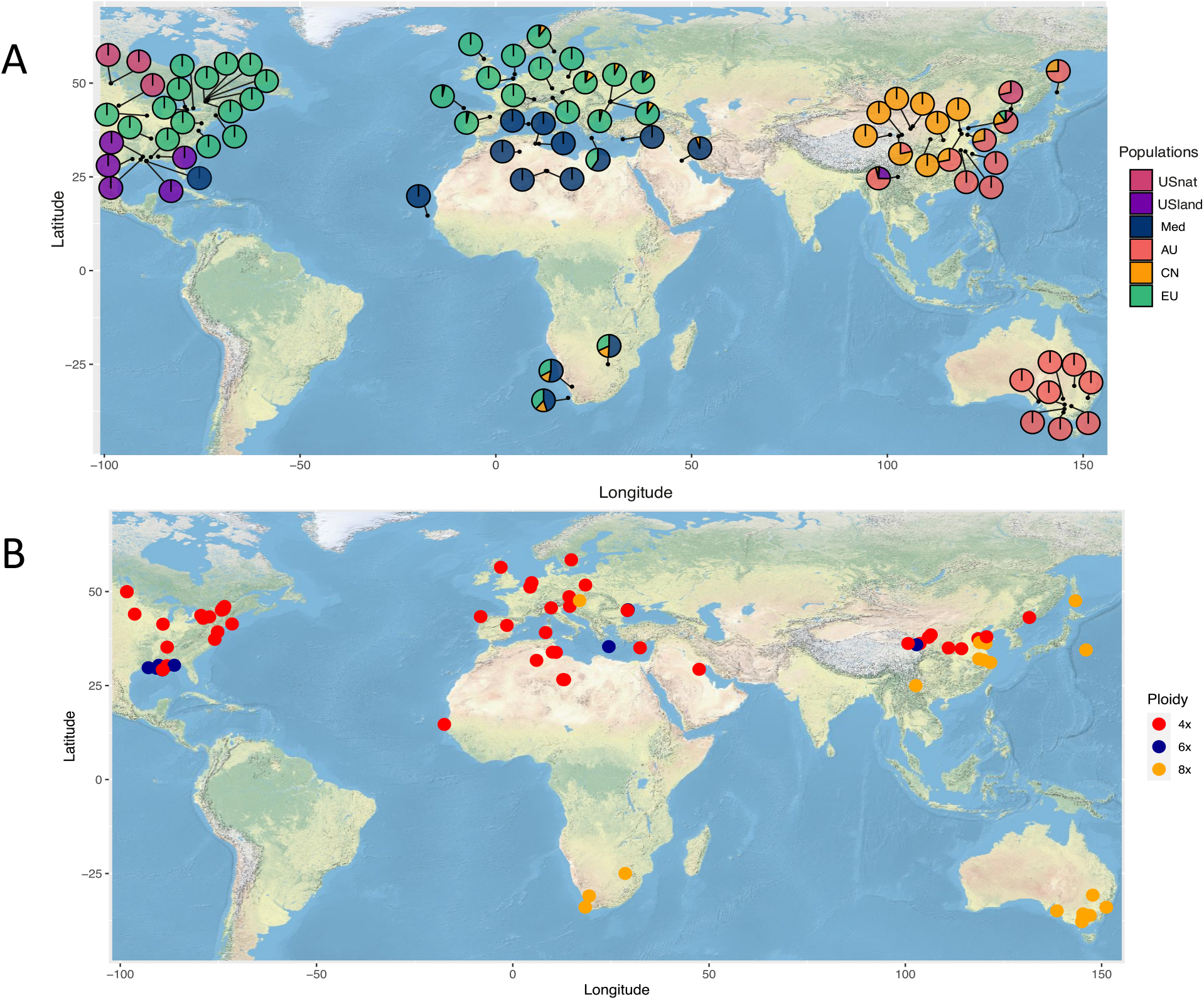
Sampling locations of *Phragmites australis* individuals in the study. A) Pie charts indicate the genetic admixture proportions. B) Ploidy level estimates from flow cytometry. The ploidy levels of the individuals with no flow cytometry data were predicted using sequencing data, see Methods for a detailed description. Different colors represent distinct ploidy level.

Based on the initial analysis, we identified three non-admixed representatives of distinct genetic lineages; the so-called US land type occurring in Gulf Coast (USland; individual Y7)(Lambertini, Mendelssohn, et al. 2012; Meyerson, et al. 2012), native US lineage limited to the North (Lambertini, et al. 2006) (USnat; Y17) and Mediterranean lineage (Med; Y21)(Lambertini, et al. 2006). The three accessions were sequenced to 90x coverage using Novaseq short read sequencing and assembled using MaSurCA genome assembler after which contigs representing haplotype variants of the same locus were removed. The final assembly sizes were 922 Mb for US native lineage (Y17) and 815 Mb for Mediterranean lineage (Y21), nearly twice the monoploid Cx value (0.490, 479.22 Mbp)(Pyšek, et al. 2018). In contrast, the assembly size for USland (Y7) was 1.2 Gb. Quality assessment using universally conserved single-copy genes (BUSCO v5.2.1) yielded high values (92.3-98.0% completeness, Table 2), indicating that the draft assemblies were of comparable quality and that they had captured the gene coding space with high accuracy. We further noticed equal proportions of duplicated genes across all assemblies (34-48.9%). The duplicated genes showed significant overlap among the three assemblies (Fig. 2A, Table 2), suggesting that the duplication is biological and likely associated with the polyploid nature of *P. australis*. Genome annotation was carried out next, by combining homology information from two Poales species, *ab initio* gene predictors and publicly available RNAseq data into consensus predictions using Evidence Modeler (Haas, et al. 2008; Salojärvi, et al. 2017). The highest number of protein coding genes were predicted in USland (Table 2).

**Figure 2.**
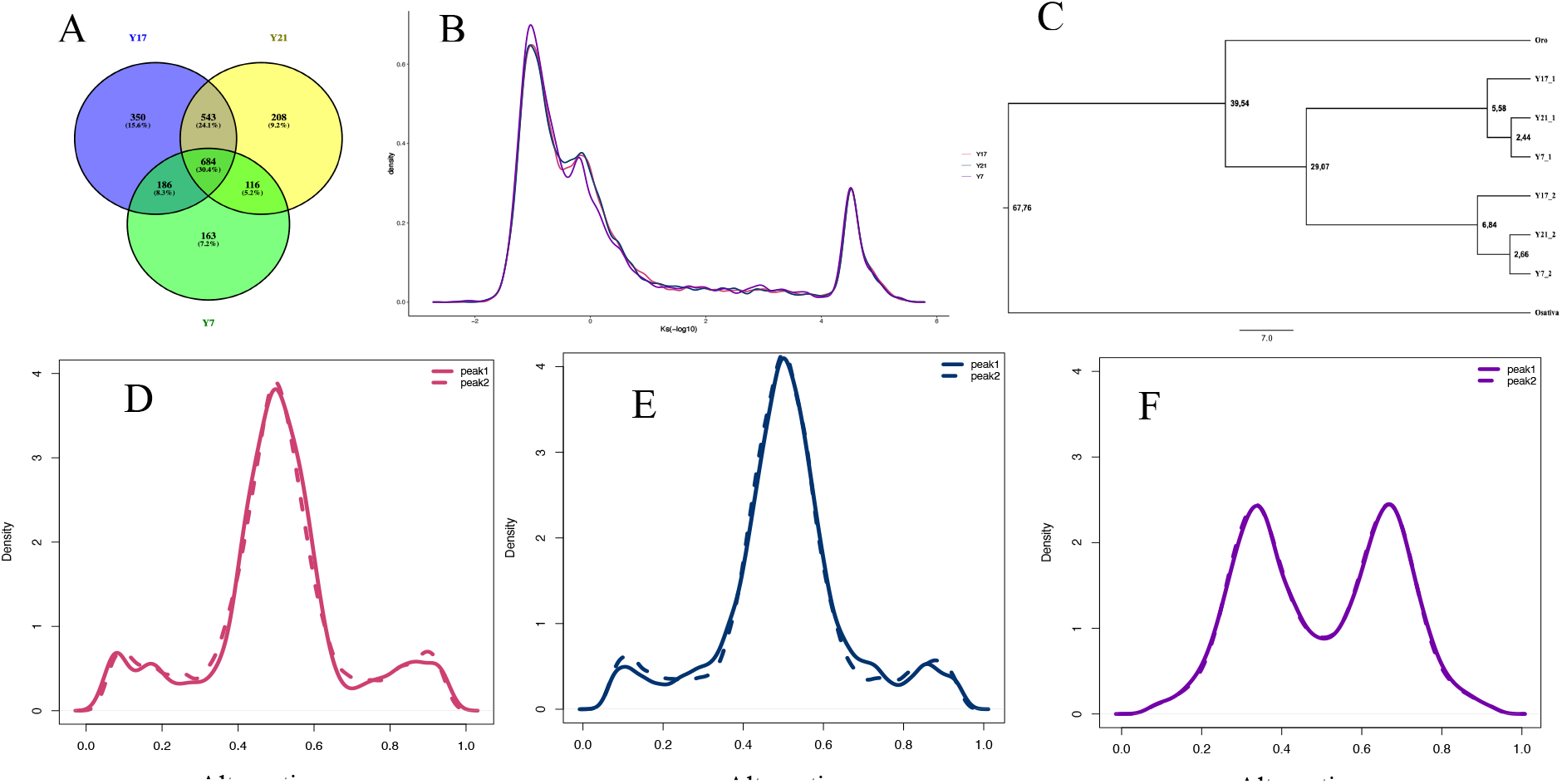
A) Overlap of the multicopy BUSCOs among genomes Y17, Y21, and Y7. B) Histograms of Ks values of syntenic genes of Y17, Y21 and Y7 genomes when aligned against *Oropetium thomaeum* genome. C) Divergence time of *Phragmites australis* subgenomes estimated from single copy genes in the ancestors and duplicated genes in current *P. australis* lineages. D-F) Allele frequency of reads mapped to peak1 and peak 2 in Y17, Y21 and Y7.

*Phragmites australis* has previously been suggested to be an allopolyploid species (Raicu, et al. 1972) but this hypothesis has been largely left unexplored due to limited evidential support. To study this hypothesis, we carried out syntenic alignments against *Oropetium thomaeum* genome assembly. For all assemblies two peaks in the synonymous (Ks) mutation spectra were detected, suggesting that the two subgenomes have different divergence times from *Oropetium*, thus confirming an allopolyploid origin for *P. australis* (Fig 2 B). All assemblies had roughly equal numbers of syntenic blocks, providing yet no explanation to the larger assembly size in Y7 (Table S1).

We next used the Ks (synonymous substitution rate) values to phase syntenic blocks with Ks values between 0 - 0.63 to subgenome A, and blocks with Ks values between 0.63 - 2 to subgenome B (Table S1). Divergence time estimation for the phased duplicates of universally conserved single-copy genes (BUSCOs), using divergence time with *O. sativa* as calibration point, showed that the subgenomes diverged at ca. 29.07 Mya (95% Highest Posterior Density, HPD 26.74 – 31.70 MYA) (Fig. 2C). Both subgenomes gave largely concordant evidence about the later divergence between the three lineages. In subgenome A, USnat lineage diverged from others at 5.58 MYA (95% HPD 4.83 - 6.40MYA), followed by divergence between Med and USland lineages at 2.44 MYA (95% HPD 2.02 - 2.87MYA). In subgenome B, USnat diverged from others at 6.84 MYA (95% HPD 6.02 - 7.72MYA), followed by divergence between USland and Med at 2.66 MYA (95% HPD 2.28 – 3.03Mya) (Fig.2 C), indicating roughly similar evolutionary rates for the two subgenomes. It is not known when the allopolyploidization event occurred, but the clear separation of the two subgenomes suggests that the two ancestral species have evolved independently for a considerable time to produce barriers for gene flow between the two subgenomes. It is worth noticing that the divergence time between *Arundo* and *Phragmites* species was also estimated to be 29 Mya(Hardion, et al. 2017), therefore one possible scenario is that the current genome is generated by ancestor of *Phragmites* hybridizing with an ancestral species of the *Arundo* genus.

### Alternative allele percentage spectrum predicts differing ploidy levels

The percent of reads supporting reference vs. alternative allele in a locus can be used as a tool to assess the ploidy levels of the individuals (Augusto Corrêa dos Santos, et al. 2017). For all individuals with whole genome sequencing data the allele percentage histograms showed a main peak at 0.5, but the overall shape differed. In USnat and Med, two side peaks were observed at 0.09 and 0.91, whereas in USland the major peak was wider and less steep (Fig. 2D-F). We hypothesized that the inconclusive shape was due to repeat elements and other ambiguously mapping regions in the genomes, and therefore focused the analysis on the gene models with assignations to subgenome A or B based on the Ks values. Both USnat and Med displayed a clear main peak at 0.5 and retained two side peaks at 0.09 and 0.91 (Fig. 2D-E), indicating the subgenomes were diploid, and thus these two lineages should be regarded as allotetraploid. On the other hand, for USland two major peaks at 0.337 and 0.669 were observed (Fig. 2F), suggesting a hexaploid genome organization; higher ploidy level also explains its 1.5x larger assembly size. Moreover, the number of conserved single-copy genes (BUSCOs) with three duplicates is much higher in Y7 (106 copies) than in Y17 (51 copies) and Y21 (58 copies), which further supports the hypothesis (Table 2). Therefore, USland lineage most likely resulted from a tetraploid – octoploid hybridization event.

Although both USnat and Med lineages were allotetraploid, the genome size and gene count of USnat was roughly 14% higher than Med lineage. The two lineages diverged at 5.58 – 6.84 Mya in late Miocene, suggesting that different lineages may have been going through differential gene loss in diploidization (Jones and Pašakinskienė 2005). There were 1916 duplicated BUSCOs shared by USnat and Med lineage, but only 1203 (62.79%) of these were also shared with USland with 1665 duplicated BUSCOs. One possible reason for the increased amount of copies in USland could be that the remaining duplicates were inherited from another closely related species or an allopatric diverged population.

### Population genetics analysis reveals links between polyploidization and lineage divergence

The initial *de novo* assembly of RAD-tags generated 4,565,536 loci with the average length of 521.3 bp and mean effective coverage of 39.0x per sample (with minimum 22.8x and maximum at 65.7x). Altogether the regions contained 40,507,058 single nucleotide polymorphism (SNP) sites. In contrast, a reference-guided alignment against USnat assembly resulted in 7,045,580 loci and 19,535,449 SNP positions, suggesting that many of the *de novo* loci were collapsed assemblies of the two subgenomes. In the reference-based RAD analysis the region length was 305.9, and the mean coverage of each sample was 39.6x (minimum coverage 23.9x, maximum coverage 65.8x). From this set, 16,500 SNPs mapped into chloroplast genome.

#### Population structure shows a split into six lineages according to geographic location

After selecting one random SNP from each Stacks locus, we obtained 538,761 variant sites in total, and 13,536 variants remained after selecting loci with no more than 10% missing values and minor allele frequency greater than 5%. Maximum likelihood phylogenetic trees revealed seven distinct genetic clusters, namely Native American lineage (USnat), Australian lineage (AU), Chinese lineage (CN), European lineage (EU), South African lineage, Mediterranean lineage (Med), and US land type (USland) (Fig. 3B). USnat and AU lineages diverged from other lineages at an early stage. Subsequently, CN lineage split from EU, South Africa, Med and USland. Population structure analyses using fastSTRUCTURE supported K=6 as the best number of clusters, in terms of highest marginal likelihood. These six genetic groups coincided with the geographical area, corresponding to USnat, USland, Med, AU, CN, and EU in the phylogenetic tree (Fig. 3B). Admixed individuals were mainly detected between AU vs CN, EU vs CN, and Med vs EU vs CN (Table 3).

**Figure 3.**
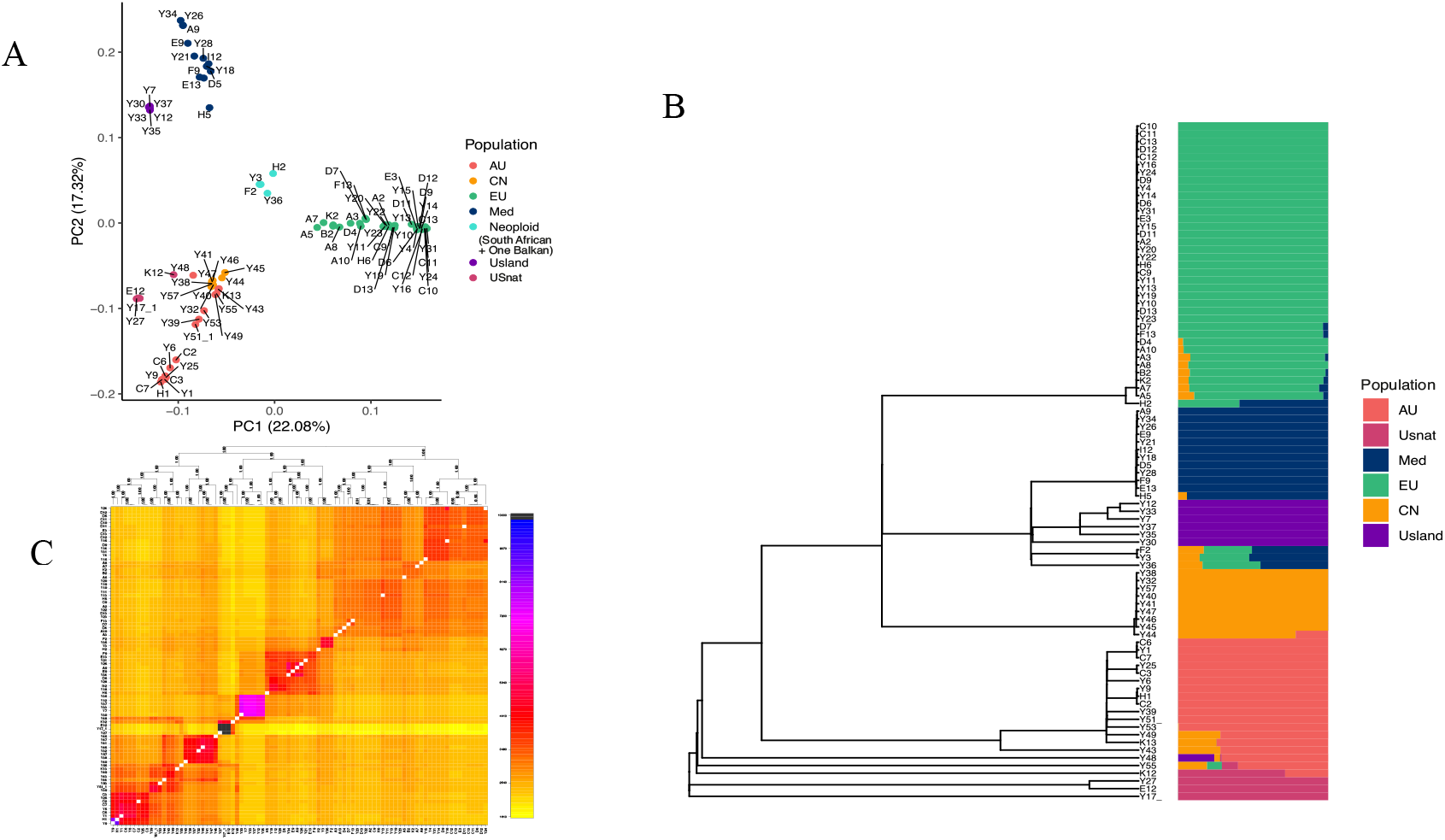
Nuclear phylogeny and population structure of *Phragmites australis* lineages. A) Population structure of *P. australis* samples from principal component analysis based on variants called against the reference genome Y17. B) Left: Maximum likelihood phylogenetic tree estimated from 3,023,618 variants aligned to Y17, with 10% of missing data allowed; Right: fastStructure population assignments based on variants. C) Genetic clusters revealed by fineRADstructure based on genetic similarities among individuals from *de novo* analysis. Darker colors indicate higher level of similarity.

The phylogeny based on RADseq loci mapping on chloroplast showed a similar split into six genetic groups corresponding to USnat, AU, USland, Med, CN and EU (Fig. 4), with varying bootstrap support values. Around 14 individuals showed discordant placement between chloroplast and nuclear trees, which is very likely a result from different origins of maternal lineages, pointing towards several independent hybridization events and extensive gene flow. For example, five individuals of Med lineage carry chloroplast from EU lineage, whereas two individuals from EU lineage and three individuals from Med lineage carry CN chloroplast; another four genetically admixed individuals carry the chloroplast of the group intermediate between main genetic groups (Fig. 4).

**Figure 4.**
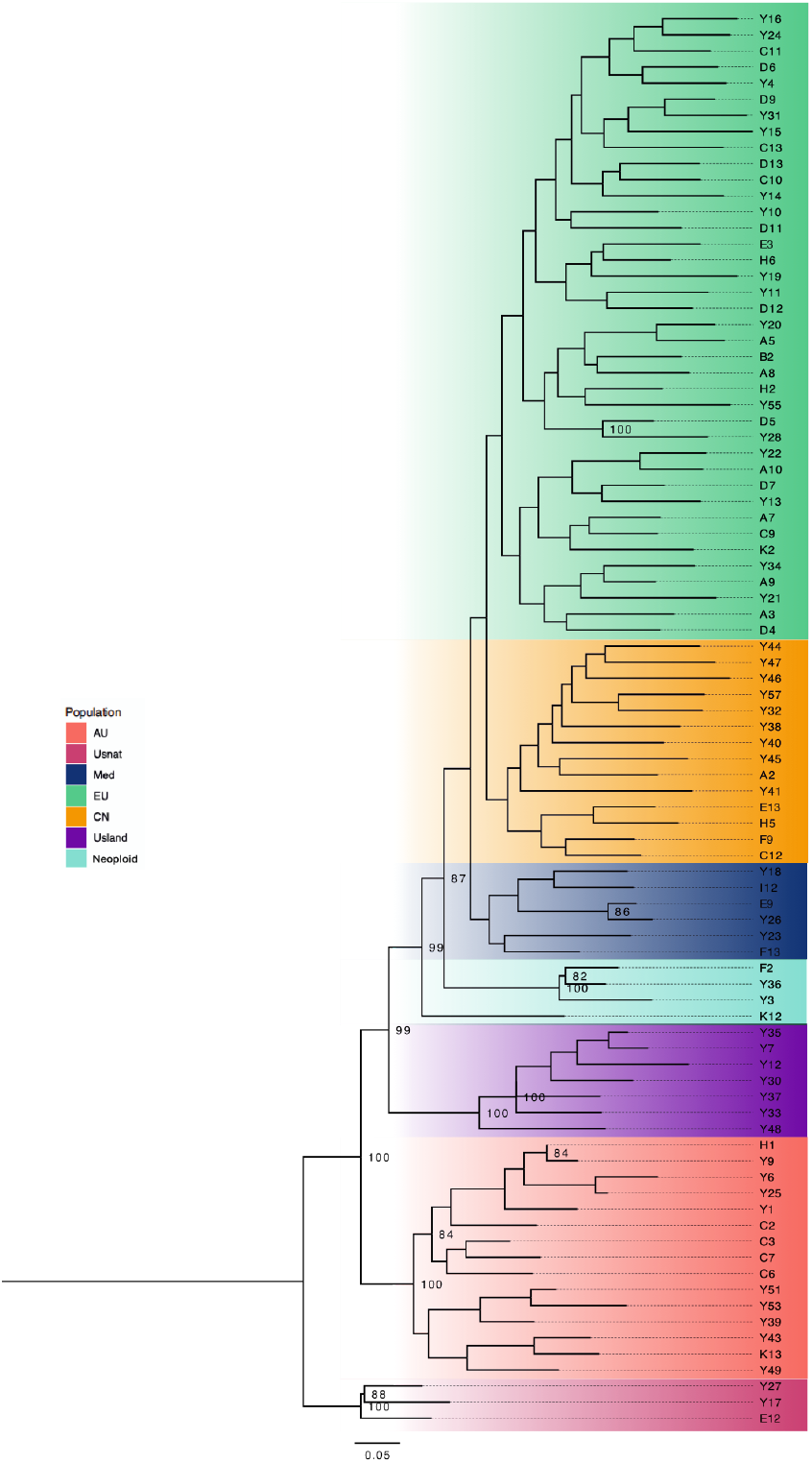
Phylogenetic tree constructed from chloroplast reads subtracted from RAD-seq dataset and mapped to chloroplast genome. Branch colors indicate the genetic groups corresponding to nuclear trees. Number on the node shows bootstrap value from 100 replicates, and only bootstrap value higher than 80 are showed.

Our data showed a rather clear geographic distribution of each genetic group: USnat lineage is distributed at the border of United States and Canada, whereas USland type is distributed at the Gulf of Mexico; Med lineage are found in Mediterranean regions, North Africa and Gulf of Mexico; CN lineage are mainly located along the Yellow River in China; AU lineage is colonizing Australia, Southern China, Pacific peninsula and islands, Korean peninsula; EU lineage is dominant across Europe, and also the invasive lineage in United States; finally, the Neoploid (South Africa) lineage is distributed in South Africa (Fig. 1). Although we could not obtain the sequences of *trn*T-*trn*L and *rbc*L–*psa*I regions to directly compare the results with previous studies based on chloroplast haplotypes (Saltonstall 2002; Saltonstall 2003; An, et al. 2012), we could infer the corresponding haplotypes by the corresponding geographic locations (Table S2). For example, the chloroplast haplotype P which is distributed in Eastern China may represent AU lineage, as they have overlapping geographic occurrences (Lambertini, et al. 2020; Liu, et al. 2020).

The geographic distribution of *P. australis* lineages was not precisely concordant with their origins, most likely due to extensive gene flow between the populations or frequent artificial transfer. For example, the nuclear phylogeny showed a subset of North American samples clustering together with a subclade of EU, indicating an artificial introduction from EU to North America (Saltonstall 2002). In addition, some introduced individuals in US (sample information obtained from H. B.) were found in several subclusters of the EU lineage suggesting multiple introductions to North America. These individuals spread quickly and are known to be invasive in North America (Saltonstall 2003).

#### Multiple ploidy levels in P. australis

For RAD-seq data, a histogram of the proportion of reads supporting alternative allele calls showed modes at around 50% in flow cytometry-confirmed tetraploids, at 0.35, 0.65 in hexaploids, and between 0.35 – 0.65 in octoploids (Fig. S1). Of the total of 88 *Phragmites* individuals, the ploidy levels of 54 individuals were quantified with flow cytometry, and only four samples showed different patterns between flow cytometry and our prediction method (Y12, Y24, Y37, K2); this could be an error in flow cytometry or aneuploidy of the same individual. Therefore, the prediction accuracy is at least 92.6% (Table 1), indicating that the prediction of ploidy levels from RAD-seq data can be done accurately. Our approach is intuitive to visualize and is suited for large plant genomes with high ploidy levels. However, it should not be treated as a replacement of traditional methods for determining ploidy level but could be used as a complementary method in addition to the flow cytometry results.

Using our approach, we predicted the ploidy levels for the remaining 34 individuals. Most individuals from North America, Northern China, Mediterranean regions, and Europe (representing USnat, CN, Med, EU lineages; see Fig. 3) were predicted to be tetraploid, showing unimodal distribution peaking at 0.5 (Table 1; Fig 5A-D). All representatives from Gulf Coast (USland lineage, Fig. 3), and a few admixed individuals (Fig. 3), were predicted to be hexaploids (Fig. 5 E-F), whereas all Australian (AU lineage, Fig. 3) and a few admixed individuals from South Africa and Mediterranean region (Fig. 3) were predicted to be octoploids (Table 1, Table 3; Fig. 5 F-J). In sum, each lineage detected in population structure analyses was characterized by its own ploidy level: The AU is octoploid, the USland is hexaploid, and the Neoploid (South Africa) is octoploid. The other lineages such as CN, Med, and most EU individuals are tetraploid.

**Figure 5.**
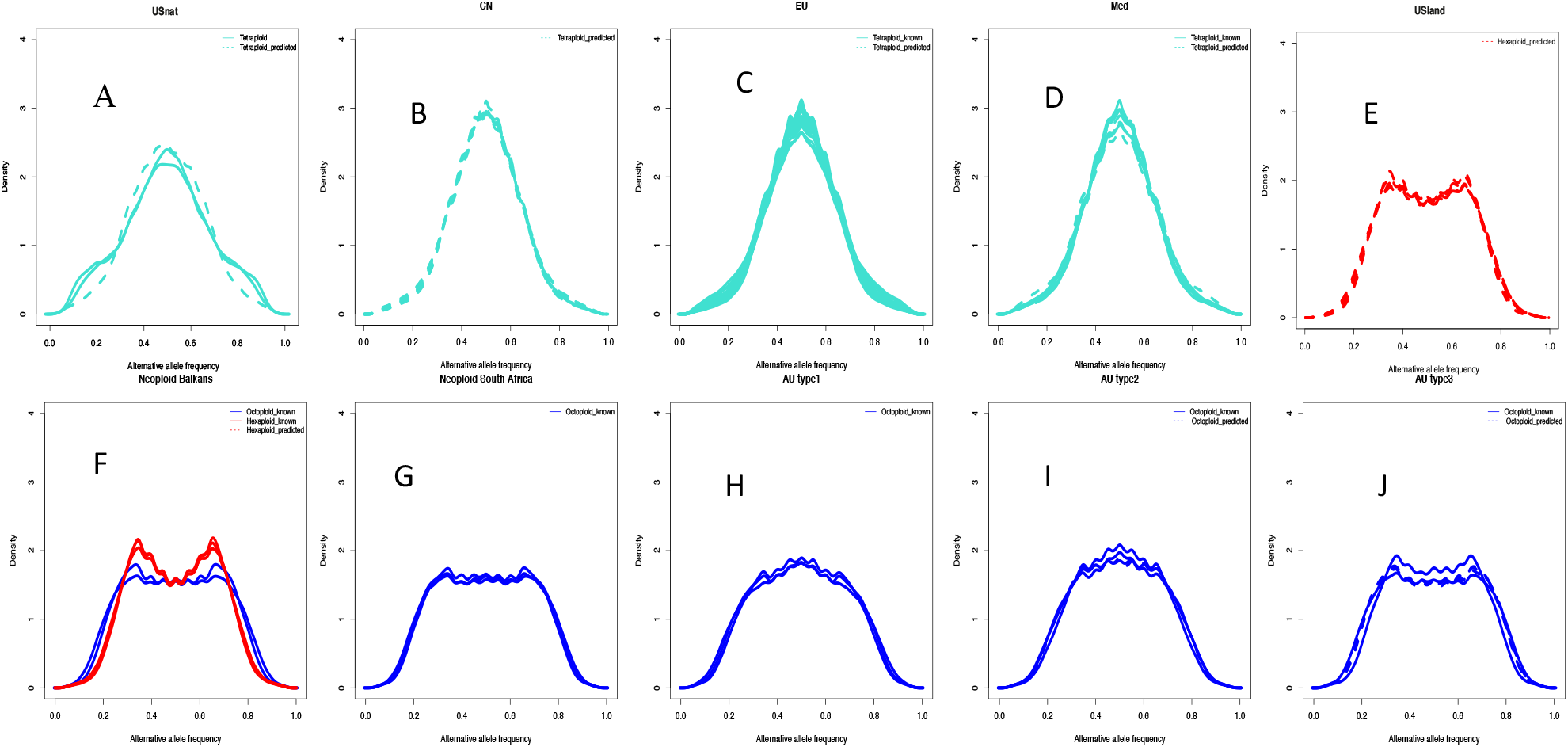
The alternative allele frequency histograms from aligning the RAD-seq reads to the Y17 reference genome. Solid line shows frequency distribution of alternative alleles in individuals where the result was confirmed with flow cytometry. Dashed line shows frequency distribution of alternative alleles in individuals with missing or differing flow cytometry measurement. Turquoise, red and blue color represented tetraploid, hexaploidy and octoploid respectively. Pattern of alternative allele frequency in population USnat (A), CN (B), EU (C), Med (D), USland (E), Neoploid (F-G), and AU (H-J).

Out of 20 admixed individuals, 13 showed increased ploidy levels and two individuals showed decreased ploidy levels compared to parental populations. Five individuals, including samples from South Africa, Romania and Hungary, were admixed offspring of tetraploid lineages CN, EU and Med (Fig. 3), but they were found to be octoploid, while another three individuals of the same background were predicted to be hexaploids, and we refer to these individuals as Neoploid (Balkans); see Fig 5F-G, samples Y3, Y36, A5, A7, A8, B2, F2, H2 and K2 (Table 3). The octoploids in the Danube delta were classified into EU lineage previously (Lambertini, Eller, et al. 2012).

#### Ploidy levels and geography explain the population structure

To test whether the different ploidy levels are visible in the SNP patterns we next assessed the population structure in *P. australis* with principal component analysis. The first three principal components explained 52.1% of the data and showed segregation pattern according to geography and ploidy level. The hexaploids largely fell between the tetraploid and octoploid population groups (Fig. 3A). The EU, USland and South African octoploids were separated from the others by the first PCA axis, and the second axis further distinguished Med and AU/CN lineages (Fig. 3A). Although AU and CN were of different ploidy levels, they were not clearly distinguishable from the PCA plot. One rare individual, Y48, with admixture from AU, USland and CN, was located between the AU and USland populations, in agreement with the admixture result (Fig. 3A).

For more accurate quantification of the covariates contributing to genetic variation we carried out redundancy analysis (RDA; Fig. 6)(Israels 1984; Capblancq, et al. 2018). Altogether 42.9% of the total genetic variation was explained by the population groups, whereas ploidy levels accounted for 9.7% of the variation. Geographic coordinates accounted for a small but significant proportion of 2.1% of variance (latitude, *p*= 0.01; longitude, *p*= 0.01, respectively) after modeling the effect of population group as covariate. The ploidy levels were confounded within the population groups, and thus the ploidy did not explain any additional variance after including the groups as covariate. Similarly, more fine-scale genetic coancestry estimation with fineRADstructure also revealed seven clusters (Fig. 4C), with most of the groups corresponding to the geographic locations, showing high genetic coancestry of AU, USland, Med, South African, USnat, CN and EU lineages. Because the South African individuals formed a group of neopolyploids with similar admixture from the tetraploid CN, EU and Med lineages, we refer the lineage as Neoploid (South Africa) in further analyses.

**Figure 6.**
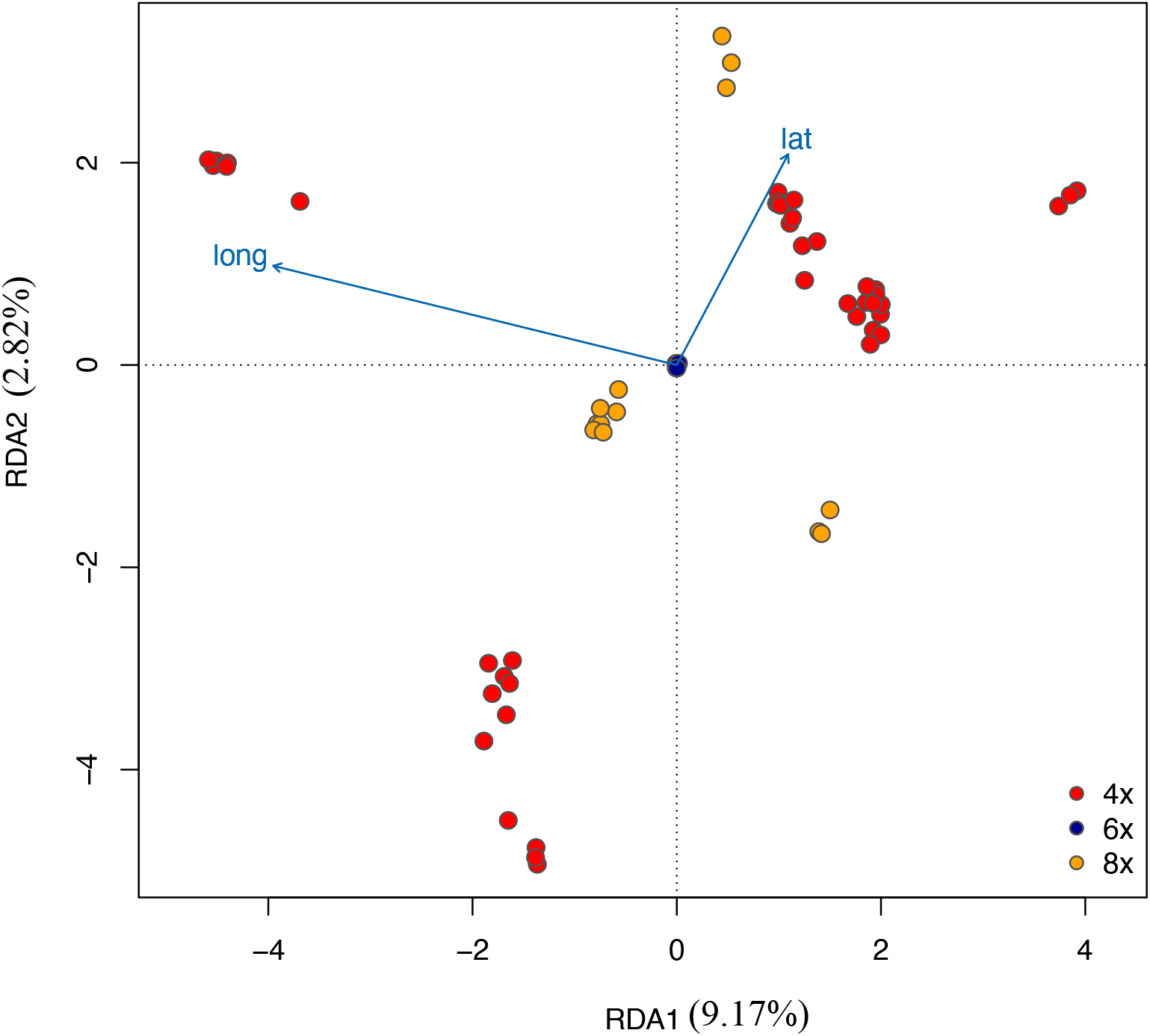
RDA analysis under constraint model using ploidy level as covariates. RDA1 explains 9.17% of the data, and RDA2 explains 2.82% of the data. The arrows lat and long indicated the latitude and longitude of sampling sites of each individual. Ploidy level explains 15.78% of the variance and latitude and longitude explains 7.87% of variance.

We next carried out modeling of population divergence times using Approximate Bayesian Computations (ABC) modeling in Fastsimcoal2 by fitting joint site frequency spectrum (SFS) two-population split models. To summarize the splits, we constructed a neighbor network based on the pairwise divergences. The topology of the network slightly differs from the phylogeny, possibly because the method is not limited to bifurcating trees (Fig. 7C), and provides a viable hypothesis that takes into account also the gene flow between populations. Altogether, a split between Asian and European populations is detected, with the hexaploid hybrid USland intemediate to Med and USnat. MRCA of USnat and the remaining lineages fell in the range 3.42 - 9.16 Mya, using generation time range of 2 - 4 years (Table 9). Moreover, AU diverged from CN at 1.74 - 3.48 Mya, and EU diverged from Med at 1.64 - 3.28 Mya, and Med diverged from USland at 1.34 - 2.68Mya (Fig. 7 C; Table 9). These populations diverged within the last 1.34 - 3.48 million years in Pliocene and Pleistocene, indicating that the climate change during Pleistocene glaciation cycles may have enhanced lineage divergence and polyploidization.

**Figure 7.**
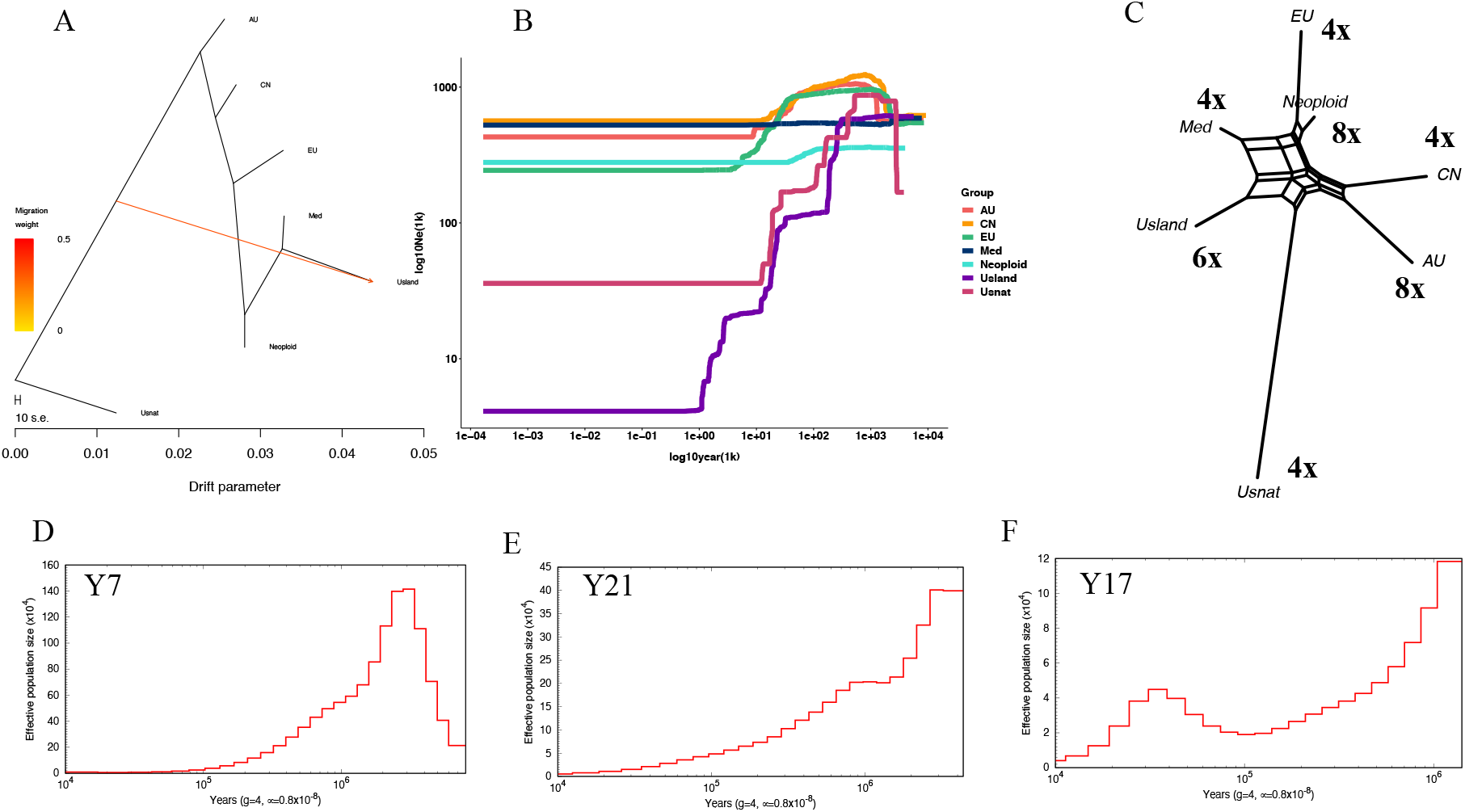
A) Migration events inferred with Treemix. B) Historical effective population sizes of seven *P. australis* lineages estimated with Stairway plot (AU, CN, EU, Med, Neoploid, USland, and USnat). Admixed individuals were excluded from the population analyses, except Neoploid. C) A neighbor net estimated from pairwise divergence times inferred with fastsimcoal2, the numbers indicate the ploidy levels. D-F) Population histories for Y7, Y21 and Y17 inferred using PSMC with mutation rate of 6.0277e–09 substitutions per site per year and generation time of 4 years.

According to the leading-edge hypothesis, source populations would preserve more genetic diversity than populations in the new territories during the spread of the species (Hewitt 1996), and studies on human populations have displayed the so-called serial founder effect, in which the diversity of the populations decreases in proportion to the distance to Africa, due to a series of founding events from diminishing populations (DeGiorgio, et al. 2009). To test whether the spread routes of *Phragmites* populations could be identified in a similar manner, we compared geographically isolated individuals from the same genetic group. For pure EU lineage, nucleotide diversity of the North American populations originating from Europe (0.12726, n = 14) is lower than within European populations (0.17707, n = 16), in support of the known founding event. For AU lineage, the nucleotide diversity of Southeastern China population (0.16872, n = 6) was higher than Australian populations (0.13283, n = 8). However, if only pure individuals of AU lineage were considered, the nucleotide diversity of Chinese population (0.13337, n = 3) was lower than the Australian population (0.16872, n = 6), illustrating the effect of admixture on nucleotide diversity. Tetraploid lineages of *P. australis* showed a decrease of genetic diversity from CN (0.096761, n=8) to EU (0.095245, n=26), Med (0.074062, n=11) and USnat (0.0207, n=3), suggesting the core lineage of *P. australis* originated in temperate grasslands of the former Laurasia (see (Pound, et al. 2011) for a reconstruction of a late Miocene vegetation map), split into Asian and European populations and further into Mediterranean population. Based on our phylogenies USnat lineage diverged early from the main Laurasian population, most likely already during the Tortonian period.

#### Extensive gene flow in Phragmites australis

Genetic divergence between USnat and any other lineage was very high (F*_st_* > 0.44, Table 8), especially with USland, see (Lambertini, Mendelssohn, et al. 2012; Liu, et al. 2018), confirming the distant genetic relationship with the rest of the lineages (Table 8). USland showed exceptionally high divergence with Neoploid (F*_st_* = 0.44689), but moderate divergence with AU, EU, CN and Med (0.29 < F*_st_*< 0.35, Table 8). The lowest F*_st_* value was observed between Neoploid (South Africa) and EU lineages (0.17126), suggesting recent gene flow (Table 5, 8). We therefore analyzed the gene flow using formal tests of introgression.

Population migrations estimated with Treemix yielded a concordant topology with the maximum likelihood phylogeny tree, suggesting one early migration event to USland lineage from an ancestral lineage not related to USnat (Fig. 7A) as the best model. However, F3 tests of admixture yielded positive values for all seven groups, providing no evidence of hybridization (Table 4). Therefore, gene flow suggested by Treemix is probably from a ghost lineage not sampled in our study (Fig. 7A). In fact, USland lineage, habituated in the Gulf Coast of North America, has been regarded as a hybrid of Mediterranean *P. australis* and *P. mauritianus* (Lambertini, Mendelssohn, et al. 2012). Similar to Mediterranean *P. australis*, the USland lineage shows significantly higher photosynthetic efficacy than EU lineage, which is probably an adaptive mechanism to the climate in its origins at tropical Africa and the Mediterranean area (Nguyen, et al. 2013). On the other hand, previous studies have showed that USland lineage holds a 200bp band in its *waxy* gene which specifically exists in *P. australis*, and also a 100bp DNA which is specific in *P. mauritianus* (Lambertini, Mendelssohn, et al. 2012), confirming its hybrid status. Hybridization between *P. australis* and *P. mauritianus* has also been observed in Southern Africa during recent decades (Canavan, et al. 2018). Due to the lack of proper outgroup for USland, we were not able to test the gene flow between USland and Med using D statistics. However, USland groups together with Med lineage in nuclear phylogenetic topologies (Fig. 4B; Fig. 7A), and the absolute genetic distance d_xy_ between USland and Med is lower than for any other lineage (Table 8), in support of the hypothesis that Med is probably the second parental lineage of USland.

For the other lineages, from the 40 quartets of lineages compared with the ABBABABA analysis using USnat as outgroup, altogether 30 quartets tested significant (Table 5), suggesting gene flow among most of the lineages (Table 5). The gene flow between the lineages was further confirmed by D statistics for randomly sampled four representative individuals from each lineage (except South Africa lineage); the individual tests confirmed the gene flow results from population-level data but also revealed possible gene flow between Med and AU, and Med and USland (Table S3). The complex pattern of admixture between nearly every population tested suggests extensive global gene flow between populations; similar conclusion can be drawn from the admixture analysis where 25% of all individuals were found to be admixed, as well as from the incongruence between nuclear and organellar phylogeny. The admixture analyses and ABBABABA tests (Fig. 2B, Table 5) further supported extensive hybridization between AU and CN lineages, suggesting the AU lineage may have been widely spread throughout the Asian continent. The CN clade was mainly confined within the Yellow River watershed, so this lineage may have occurred later from the west of China and spread along the river to eastern China.

### Hybridization and polyploid diversity

For RADseq data, nucleotide diversity (π) was estimated to be highest in octoploid AU lineage (0.099908, n=12) and lowest in USnat lineage (0.0207, n = 3) (Table 6), while other lineages showed intermediate levels of polymorphism (π ranging between 0.067431-0.096761, Table 6). On the other hand, the genome assemblies yielded low nucleotide diversities of neutral sites (π_s_) for USnat (Y17; 0.001142), Med (Y21; 0.003121), and higher for USland (Y7; 0.00947) (Table 7). Similar results were obtained for RADseq data when looking at individuals separately (Table 1). The inbreeding F coefficient was positive for all accessions, being highest among USnat lineage and lowest in USland lineage, suggesting a higher level of inbreeding in USnat and relatively more outbreeding in USland, correspondingly (Table 6). Tajima’s D value was positive in all populations, being moderately high in AU (0.75 ± 0.85), EU (0.66 ± 1.03), CN (0.575 ± 0.91), Med (0.63 ± 0.98), Neoploid (South Africa) (0.81 ± 0.77) and USnat (0.39 ± 0.94) lineages, and exceptionally high in USland lineage (1.54 ± 1) (Table 6).

Low nucleotide diversity and high Tajima’s D are in general hallmarks of a population bottleneck, and we therefore continued by assessing the population history of the species. Using Stairway plot with the site frequency spectrum from RADseq data and four years as generation time, an early population increase was detected between 1-5 Mya for USnat, AU, CN and EU lineages (Fig. 7), followed by a recent decline of effective population size in nearly all the lineages except Med (Fig. 7B). The decline of AU, CN and EU populations started from 50Kya. Population of USland dropped dramatically three times with first one at 500Kya, followed by a drop at 40Kya, and 5Kya. USnat started to decrease monotonously from 700Kya.

Using the same generation time, PSMC analysis on the genome assemblies showed population decreases in Med lineage (Y21) starting from around 3 Mya. In USland the decrease started from 1.5 Mya, and in USnat from 1 Mya, the latter demonstrating a slight increase at 35Kya followed by another decline (Fig. 7 D-F). Divergence time estimation had showed USland to diverge from Med lineage at 1.34 – 2.68 Mya (Fig. 2C, Table 9), and the PSMC showed the sharp increase of the population size of USland at around 2.5 Mya (Fig. 7E), indicating the hybridization event may have happened at a suture zone when the population went through contraction and expansion during glaciation cycles, or then the increase in effective population size merely reflects the hybridization event and resulting increase in heterozygosity. USnat on the other hand was also characterized with a population increase at around 5 Mya (Fig. 7B), coincidentally in line with expansion of North American Prairie in late Miocene, suggesting that the climate cooling and drying may have facilitated the lineage divergence(Retallack 1997). Towards holocene, the USland and Med populations went through an acute decline as seen in PSMC and Stairway plots (Fig. 7D, Fig. 7B) and also from the high Tajima’s D value (Table 6).

### Genetic diversity suggests transitions in reproductive mode

The decline seen in population histories was unexpected, since the population-level diversities were relatively high, and especially when considering the invasiveness of the expanding EU lineage. We suspected that the results might reflect a break of self-incompatibility and transition into clonal propagation, since the transit of reproductive mode strongly affects genetic diversity regarding the mutation accumulation, allele frequency alteration, and population size changes as time passes. Population genetic simulations predict that compared to outcrossing and selfing lineages, a clonal lineage experiences relaxed selection and thus accumulates more deleterious mutations resulting in an overall high level of heterozygosity. However, due to lack of recombination, genotypic diversity in asexual population would be low since genotypes which have accumulated a high number of deleterious alleles, and therefore have reduced fitness, would be purged from the population (the phenomenon is known as Muller’s ratchet). Additionally, clonal populations may undergo frequent bottlenecks, since they can colonize habitats with just a few individuals. Altogether these effects result in effective population decline, hypothesized to lead into an evolutionary dead-end in the long run (Glemin and Galtier 2012). The predictions have been verified empirically in both animals and plants, such as *Primula* species (Wang, et al. 2021), stick insects (Bast, et al. 2018), duckweed (Ho, et al. 2019), bdelloid rotifers (Barraclough, et al. 2007), and wasps (Tvedte, et al. 2020), to name a few.

*P. australis* uses both sexual and asexual reproduction mode. Genome evidence pointed towards frequent selfing among *Phragmites* populations, since the genome assemblies demonstrated low levels of heterozygosity and the RAD-seq data yielded high values of inbreeding coefficient. However, seed set rate of *P. australis* is very low (mean 9.7 −12%) due to its partial self-incompatibility (Tachibana 1984; Ishii and Kadono 2002), indicating that asexual reproduction may also be playing an important role. In fact, clonal propagation has been shown to work efficiently in *Phragmites* (Amsberry, et al. 2000; Alvarez, et al. 2005; Čuda, et al. 2021).

Relaxed selection would result in an increased ratio of non-synonymous to synonymous mutations in asexual and selfing proliferation (Henry, et al. 2012; Lovell, et al. 2017), whereas in outcrossing species purifying selection would limit the amount of deleterious alleles (Glemin and Galtier 2012). Nucleotide diversities of deleterious sites (π_n_) were Y17 (0.001417), Y21 (0.002784), Y7 (0.008596) (Table 7), and the corresponding ratios of non-synonymous to synonymous diversity, π_n_/π_s_, were 1.24 for USnat, 0.89 and 0.91 for Med and USland, respectively. The ratio is very high compared to other plants (Chen, et al. 2017) and suggests relaxed to no selection on deleterious sites (Johnson and Howard 2007). The USnat lineage had the highest ratio of nonsynonymous π_n_ to synonymous π_s_ (Table 7). The USnat population has long been isolated from other *Phragmites* lineages, which was also reflected in the high F_*st*_ statistic (Table 8). The site-wise genetic diversity in USnat population is rather high 0.0207 (Table 6) but the heterozygosity of each individual quite low, suggesting the individuals have accumulated mainly private mutations, complying with the predictions of clonal lineage, and these alleles may have fixed into homozygosity due to selfing. The overall diversity is therefore possibly a result of a combination of clonal propagation (high π_n_, high population level π_s_), inbreeding (low individual level π_s_), and little gene flow from other *Phragmites* populations (increased F*_st_*). Therefore, we infer that the secluded USnat lineage may rely even more on clonal propagation than the other lineages. In fact, it has been found in North America that while the long-distance dispersal of USnat is facilitated by seedlings, the local growth is mainly proceeding through vegetative propagation (Albert, et al. 2015). The combination of selfing and clonal propagation possibly also explains the sharp decline of effective population size of Usnat since 700Kya (Fig. 7B).

## Discussion

### Intraspecific hybrids give rise to new polyploids at contact zones

Altogether, genetic admixture was commonly seen between the borders of lineages, suggesting low reproductive barriers among lineages. Neoploid lineages occurring in South Africa and Balkans are derived from admixture of the tetraploid Med, EU and CN lineages (Fig. 3, Table 3). The neopolyploids are special in the sense that all the parental lineages are tetraploid, but the resulting offspring are either hexaploids or octoploids. Three out of four individuals from South Africa are admixed by Med, EU and CN tetraploid lineages. All the three samples, located far from each other, were predicted to be octoploids (Fig. 1), concordant with earlier records (Clevering and Lissner 1999).

Admixed octoploid individuals were found also in Hungary and Romania, with a slightly different allele frequency pattern from South African populations (Fig. 5 F-G), suggesting this may be an independent polyploidization event caused by intraspecies genomic conflicts. Not all admixed individuals had their chromosomes doubled (Table 3). In fact, only the individuals simultaneously carrying alleles of Med, EU and CN ancestry had higher ploidy levels. These individuals group within EU lineage in both chloroplast and RAD-seq genomic phylogeny, implying recursive hybridization with EU lineage after polyploidization. Beyond this study, other researchers have found octoploids in countries around Mediterranean Sea, including Romania, Greece, Afganistan, Iran, Algeria, Morocco, Tunisia, France, Spain and previous Yugoslavian area (Gorenflot 1986; Connor, et al. 1998). These regions overlap with the borders of Med and EU lineage, possibly also the CN lineage, and thus hybridization and polyploidization events are very likely to happen frequently there. Concordantly, intensive investigation has shown the existence of a mixture of octoploids, tetraploids and hexaploids in the Danube Delta, likewise inferred to originate from recent polyploidization and hybridization (Raicu, et al. 1972; Lambertini, Eller, et al. 2012). The hexaploids in Greece and Romania were found to be admixed individuals with tetraploid ancestors. This is supported from previous studies showing seeds from the same inflorescence producing offspring of different ploidy levels, suggesting the hexaploidy may be generated by hybridization of tetraploid and octoploid (Paucã-Comãnescu, et al. 1999). The same circumstance occurred in *Spartina*, where the dodecaploid *S. anglica* resulted from whole genome duplication after the hybridization of two hexaploid species *S. alterniflora* and *S. maritima*, followed by a recurrent hybridization with coexisting parental species *S. alterniflora*, giving rise to nonaploid individuals (Renny-Byfield, et al. 2010).

### Asexual propagation may contribute to genomic conflict

Autopolyploidization and allopolyploidization have been regarded as the main ways of polyploidization, with the underlying mechanism of how whole genome duplication happens still being largely unknown (Stuessy and Weiss-Schneeweiss 2019). If the parental species are well diverged, the genomic incompatibility after hybridization may give more chance of mistakes in meiosis, thus leading to new polyploids. In this study, neopolyploids were inferred to result as admixture from three lineages of the same species, implying that the regulation of genomic networks may be broken down. One possibility is frequent misregulation due to stressful environmental conditions. Extreme environment has been observed as one way to induce polyploidization, especially autopolyploids. One example is the *Brachypodium distachyon* in Iberian Penisula, where the distribution of diploids and allotetraploids was significantly associated with the dryness of living environment; arid environment may cause problems with cytotype segregation and thus enable the polyploidization in Poaceae species (Manzaneda, et al. 2012). In *P. australis*, the occurrence of octoploids is mainly distributed in Australia, South Africa, or between Middle Europe and Black Sea, where temperature is high throughout the year. Thus, the highly stressing, dry and hot climate may have caused errors in meiosis resulting in unreduced gametes, and the resulting plasticity may have enhanced their ability to adapt to the new ecological niche (Manzaneda, et al. 2012; Stuessy and Weiss-Schneeweiss 2019). However, this conclusion is controversial since an experimental study sampling *Brachypodium* species along the aridity gradient in Israel denied the hypothesis that dryness increases the chance of allopolyploidization (Achenbach, et al. 2012).

Since these polyploids were observed mainly along the contact zones, a possible genomic component contributing to the breakup follows from the extensive clonal propagation in *Phragmites* and the resulting relaxed selection and accumulation of deleterious alleles. Long term accumulation in allopatric populations may have led into increased genomic divergence such that intraspecific genomic conflicts arise among progeny from these diverged populations. This could be due to misregulation during meiosis; since sexual propagation is of lower importance in a predominantly clonally propagating species, the genes associated with it are not under strict purifying selection. Taken together, we suggest that decreasing population sizes in *Phragmites* first led into adopting selfing as propagation strategy. This resulted in further decline of population sizes and perhaps widespread inbreeding depression, and therefore clonal propagation arose as a solution. The accumulation of deleterious alleles then eventually results in genome incompatibility and polyploid hybrids are formed when long diverged populations encounter at contact zones.

## Materials and Methods

### DNA extraction, RAD sequencing and whole genome sequencing

A total of 88 *P. australis* individuals were obtained from Eurasian, North American, Oceanian and African continents (Fig. 1, Table 1) and planted in Aarhus University and Shandong University. Some of the individuals have already been used in previous phylogeographic studies, and thus we were able to link them to the defined lineages in other studies (see more in Table 1). One *Arundo donax* (giant reed) individual was included as outgroup. For RAD sequencing, DNA was extracted from fresh leaves using CTAB method. The leaves were ground into powder and mixed with 500 μl extraction buffer (containing 100mM Tris pH8, 1.4 M NaCl, 20mM EDTA, 2% CTAB) and 2%β-mercaptoethanol. After incubation at 65 °C for 10 minutes, the suspension was mixed with 500 μl of chloroform and spun at 10000 rpm for 5min. Then, 500 μl of chloroform was added to the supernatant and spun at 10000 rpm for 10 minutes, and finally 800 μl ice cold ethanol was added. The tube was left in the freezer overnight and subsequently spun at 13000 rpm at 4°C for 20 minutes, followed by washing with 500 μl 70% ethanol. The pellet was precipitated at 6000rpm at 4 °C, and let dry and dissolve in 200 μl H_2_O. Paired-end RAD-seq library preparation and sequencing was completed by Shanghai Honsunbio Limited company (http://www.honsunbio.com/), using Illumina HiSeqX10. Whole genome sequences of *Oropetium thomaeum* (genome accession: Phytozome (Goodstein, et al. 2012).), *Oryza sativa* (genome accession: Phytozome), and *Miscanthus sacchariflorus* (genome assembly accession: GCA_002993905.1, NCBI) were used as outgroups. Illumine reads of *Oropetium thomaeum* (SRR2083762), *Miscanthus sinensis* (SRR486617), *Arundo donax* (SRR4319201), *Arundo plinii* (SRR4319202) and *Sorghum bicolor* (SRR12628364) were used to construct ancestral alleles of *Phragmites* lineages.

### Draft genome assembly

Based on initial analyses with RAD-seq data, we selected three individuals (Y7, Y17, Y21) representing USland, USnat and Med lineages, respectively, to be sequenced for whole genome assemblies. Following DNA extraction with CTAB protocol, the genomes were sequenced to 90x high coverage (assuming 1 Gb genome size) by NovogeAIT at Singapore using Novaseq platform. After receiving the sequencing data, the genome size was first estimated using Kmergenie (Chikhi and Medvedev 2014). Then, after quality assessment of the reads using FastQC, the three genomes were assembled using MaSuRCA assembler (Zimin, et al. 2013) with default settings. The assembly was passed through purge haplotigs v1.0.4 (Roach, et al. 2018b) to remove the redundant haplotigs resulted from the highly heterozygous regions. The completeness of the genome assemblies was assessed using BUSCO v5 (Simão, et al. 2015). and odb_database version 10, and the quality of the assembly was estimated using Quast (Gurevich, et al. 2013). The draft genome assemblies were next filtered using Purge Haplotigs pipeline to remove heterozygous genome regions which may have been accidentally assembled into two haplotigs (Roach, et al. 2018a).

### Gene prediction

Gene prediction for the three draft genome assemblies was performed by combining homology - based method and *ab initio* methods. The draft genomes were then searched against the Poaceae grasses *O. thomaeum* and *M. sinensis* to obtain the annotation information based on homology prediction. *Ab initio* gene prediction was performed using GeneMark-ES (Lomsadze, et al. 2005), BRAKER2 (Hoff, et al. 2018), PASA (Haas, et al. 2011) and AUGUSTUS (Stanke, et al. 2008). RNAseq data of *P. australis* was obtained from NCBI (accession number: SAMN04544298), aligned to each draft assemblies by STAR aligner (Dobin, et al. 2013), and used as evidence to define intron borders in BRAKER2 prediction. In addition, the RNAseq data was also assembled *de novo* using TRINITY (Grabherr, et al. 2011) and used in PASA for generating a high quality dataset for downstream *ab initio* gene predictions. Finally, all evidence of gene prediction was combined in EvidenceModeler (Haas, et al. 2008) to get consensus gene model predictions. The completeness of predicted gene models was assessed using BUSCO by searching against poales_odb10 protein database. Significance of overlap between duplicated BUSCOs in different genome assemblies were tested with Fisher’s exact test using R package “GeneOverlap”(Shen 2014).

### Syntenic blocks and divergence of subgenomes

The three draft genomes were each aligned against *O. thomaeum* genome (Genome ID 51527 in CoGe database, https://genomeevolution.org/coge) using SynMap tool in CoGe (Lyons and Freeling 2008) that identifies syntenic blocks. The Ks (synonymous substitutions rate) rates for the syntenic genes were then downloaded from CoGe and syntenic blocks were phased into two subgenomes based on the average Ks values throughout the block.

To date the time of WGD in relation to speciation, we inspected BUSCO genes that were present in rice (*O. sativa*), *O. thomaeum*, Y7, Y17 and Y21. *Oryza sativa* and *O. thomaeum* were used as outgroups. For polyploids, multi-copy genes sourced from several polyploidizations often result in incongruent phylogeny. Single copy orthologs are better choice to reflect the species evolution due to less complicated history, and allopolyploidization events could be traced by analyzing the two copies of paralogous gene sets belonging to their own parental species (Brysting, et al. 2011). Following this idea, we used the BUSCOs which were present as a single copy in the outgroups and duplicated in Y7, Y17 and Y21 to find out the divergence time of the subgenomes. These BUSCOs were considered to be the orthologs in all species, but paralogs in *Phragmites* species, representing the two ancestral subgenomes. In total 580 such BUSCOs among *O. sativa*, O. *thomaeum* and *Phragmites* lineages were obtained. From this set, we randomly selected 25 BUSCOs which showed coherent topology with species tree and estimated divergence time of *Phragmites* subgenomes using BEAST2 with calibrated Yule model. A site model of HKY+Gamma was used, with four categories of Gamma distribution, and the molecular clock was set to be random local clock which allows certain variation of clock rate among lineages. Based on fossil record in Oryzeae and calibration point of BOP and PACMAD lineages (Prasad, et al. 2011; Christin, et al. 2014), we set the 95% confidence interval of total most recent common ancestor (tmrca) to be 65.6 −73.1Mya, by assigning a lognormal prior, with an offset 65.0, a mean of 3.0, and a standard deviation of 0.8 (Mean In Real Space). The total MCMC sampling chain length was 10 million iterations, and the states were stored every 5000 samples. The divergence estimation was run twice with the same parameter to guarantee the convergence, with each run achieving ESS value of all parameters to be higher than 200.

### Identification of putative deleterious mutations

The nucleotide diversites of neutral and deleterious sites were calculated following the procedure in https://github.com/jsalojar/PiNSiR. Briefly, the short reads of Y17, Y21 and Y7 were aligned to their own draft genome using bwa mem to obtain the variants along the genome, sorted with Samtools (Li, et al. 2009), and genome coverage variants were called using bcftools (Li, et al. 2009). Site-wise diversities were estimated using ANGSD (Korneliussen, et al. 2014) and used as input in the R package PiNSiR. The ratio of nonsynonymous/synonymous mutations of each subgenome was performed using the same method and esimtating the diversity using the coordinates of the syntenic regions with *O. thomaeum*.

### Variant calling

RAD-tag sequences were processed following Stacks pipeline refmap methods by aligning the reads to the reference genome Y17 (Catchen, et al. 2013). We first demultiplexed the reads and removed the barcode using process_radtags. In the refmap pipeline, the paired reads were aligned to Y17 assembly representing USnat lineage using Bowtie2 (Langmead and Salzberg 2012). The RAD-seq data was also aligned to the already published chloroplast assembly (accession number: KJ825856), so as to draw the genetic information carried by organelles, and to compare the evolutionary history between nuclear and chloroplast genome.

### Ploidy level prediction

Although the ploidy level of most individuals was known from flow cytometry, the knowledge of genome size of all individuals would give a clearer picture about the evolutionary path of polyploidization. To find the link between genomic alleles and ploidy levels, we developed a method that predicts ploidy level by counting the proportions of alternative alleles based on the reference guided aligned reads. After mapping the short reads to the reference genome, similarly but differently from ploidyNGS (dos Santos, et al. 2016), we count the number of alleles that are different from the reference genome, by setting the total read coverage to range from 20 to 200, only alleles with coverage higher than 7 were considered as variants. The number of chromosome copies were predicted by counting the number of peaks of alternative alleles. For example, in diploids harboring two copies of alleles, the proportion of reads supporting alternative alleles would be around 0.5 in heterozygous sites, and thus the genomic distribution of the proportion of reads supporting alternative alleles should demonstrate a single peak at 0.5. Similarly, for a tetraploid individual, the alternative allele proportion should show peaks at 0.25,0.5,0.75, and at 0.167, 0.33, 0.5, 0.67 and, 0.83 in hexaploids etc. Based on this pattern, we predicted the ploidy levels by plotting the density of alternative allele frequency, and verified the predictions using the flow cytometry data. We performed the analysis by counting the read depth of alternative alleles in bam files after aligning the filtered reads to the reference genome Y17.

### Phylogeny

For further analyses a filtered set of high quality SNPs was obtained by selecting SNPs that are present in >90% of the samples using vcftools (Danecek, et al. 2011). Phylogeny was estimated for the set of filtered SNPs using RAxML (Stamatakis 2006) under the GTRGAMMA substitution model with 100 bootstrap replicates. The final tree was annotated and viewed in Figtree (http://tree.bio.ed.ac.uk/software/figtree/). Variants from chloroplast were also used for estimating the phylogenetic tree using same methods.

### Population structure and reticulate evolution

The population structure was estimated on the whole dataset using fastSTRUCTURE with K ranging between 1…15 (Raj, et al. 2014), best K was selected by the highest marginal likelihood. To further evaluate the genetic ancestry of each independent allele, we reduce the effect of linkage disequilibrium by selecting a random SNP from each stacks loci, and removing ninety percent of the missing data were removed using vcftools v 0.1.16 (Danecek, et al. 2011), and filtered out the data with minor allele frequency less than 0.05 using Plink 1.9 (Purcell, et al. 2007). The processed data was used to performed genetic clustering using fineRADstructure (Malinsky, et al. 2018).

Principal component analysis (PCA) was calculated using Plink 1.9 (Purcell, et al. 2007) and plotted in R to reveal the ordination of the genetic data. In addition, we performed Redundancy Analysis (RDA) using R package “vegan” (Oksanen, et al. 2010) to quantify the proportion of the genetic variation explained by covariates such as different lineages, geographical location and ploidy level, with 100 permutations to assess the importance of fitted models.

To understand the genomic structure of each lineage, we grouped individuals to populations based on fastSTRUCTURE and phylogeny results (Table 4). Inbreeding *F* coefficients of each individual were calculated using Plink v1.07 (Purcell, et al. 2007), and group mean and standard deviation group were measured. Nucleotide diversity (pi), private alleles (N_A_) and genetic divergence between populations (F*_st_* and d*_xy_*) were calculated using populations program in Stacks (Rochette, et al. 2019). F coefficient for each individual was calculated using Plink 1.9 (Purcell, et al. 2007), and the nucleotide diversity (pi) of each individual was measured using vcftools v 0.1.16 with 1000bp window size (Danecek, et al. 2011). To test for admixture, we performed f3 tests using Admixtools 2 (Patterson, et al. 2012). ABBABABA were considered to be significant only when the absolute value of Z score is higher than 3. To test the potential gene flow between lineages, we selected four pure individuals from each lineage (except for USnat lineage where one individual was used as an outgroup) and performed ABBABABA statistics among all combinations using AdmixTools v7.0.1 (Patterson, et al. 2012). Tajima’s D of each population was measured using vcfkit (Cook and Andersen 2017) using non-overlapping sliding windows of size 10,000 bp to infer whether population demographics has been heterogenous along the genome.

### Population demographics in history

Site frequency spectrum (SFS) of derived alleles were used in Stairway plot (Liu and Fu 2015) to infer the historical population size changes. To further validate our inference about demographic changes, we aligned the reads of the three whole-genome-sequenced individuals to their own reference genome using bwa (Li and Durbin 2009), and performed demographic analysis using PSMC (Li and Durbin 2012). Although it usually takes 1 year for *P. australis* to sprout and mature (Phragmites resource, 1989:15), we set the generation time to 2-4 years because it is perennial. We set the mutation rate to be the Poaceae silent-site mutation rate, 6.0277e–09 substitutions per site per year (De La Torre, et al. 2017). Number of migration events between *P. australis* lineages were tested using Treemix (Pickrell and Pritchard 2012). All pure individuals belonging to the lineages were included in the analysis, and SNPs were filtered to only include the ones that appear in all populations. We analyzed 1-5 migration events, and each scenario was tested with 5 replicates. The optimal number of migration were determined using R package “optM” based on the second-order rate of change in likelihood (RRs 2019).

To determine the derived alleles in *Phragmites* species, we aligned the genome reads of *O. thomaeum*, *A. donax*, *A. plinii*, *S. bicolor* and *M. sinensis* to Y17 using bwa mem, and then obtained consensus ancestral genome using ANGSD (Korneliussen, et al. 2014). Ancestral alleles of the variants called by refmap pipeline were called and annotated using samtools and bcftools. The multidimensional unfolded Site Frequency Spectrum (SFS) for all seven lineages was calculated using easySFS program (https://github.com/isaacovercast/easySFS). We selected the optimal projection by balancing the number of segregating sites and adequate number of individuals, resulting between 933 and 2895 segregating sites per population group. The parameters with regard to population splitting were estimated with fastsimcoal version 2.6(Excoffier, et al. 2013), and mainly focused on the divergence time between populations. In each cycle, the program run for 200,000 iterations to estimate the expected SFS and conduct 40 optimization (ECM) cycles to estimate the parameters. To find the best fitting parameter, we performed 100 runs for each scenario, and selected the best run by the highest likelihood. We used a mutation rate of 6.0277E^−09^ site/year (De La Torre, et al. 2017) as a standard, assuming two to four years of generation time for *Phragmites* species. The results were visualized using neighborNet available in R package Phangorn.

## Acknowledgements

This work was supported by International Postdoctoral Exchange Fellowship Program of China Postdoctoral Science Foundation (C.W.), the National Natural Science Foundation of China (Grant No. 31770361 to WH. G, Grant No. 31800299 to T. W., and Nanyang Technological University startup grant and Academy of Finland (decisions 318288 and 319947) to J.S. We would like to thank Pasi Rastas for providing the script to calculate alternative allele count, Sitaram Rajaraman for sharing the work pipeline of genome annotation, and Carla Lambertini for the help of geographic coordinates corrections. Finally, we want to acknowledge CSC–IT Center for Science, Finland, and NTU HPCC, Singapore, for computational resources.

## Data Availability Statements

The data underlying this article are available in NCBI SRA database and can be accessed with the BioProject ID PRJNA753984.

## Author Contributions

C.W., WH.G. and J.S. conceived the study; H.B., LL.L., F.E. and T.W. maintained and collected the samples as well as provided sampling information; C.W., LL.L. and MQ.Y. performed DNA extraction; C.W. and J.S. lead the data analysis; C.W. and J.S. wrote the paper with input from WH.G., LL.L., F.E., H.B., T.W., and MQ.Y.

