## Supplemental Figure for "Genome-wide analysis tracks the emergence of intraspecific polyploids in *Phragmites australis*"

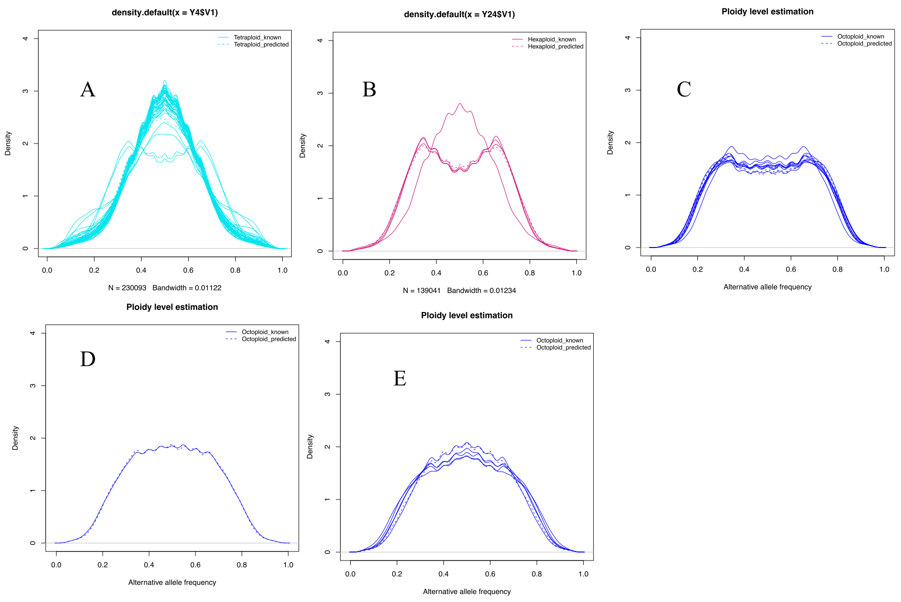


Figure S1 The alternative allele frequency histograms from aligning the RAD-seq reads to the Y17 reference genome. Solid line shows frequency distribution of alternative alleles in individuals where the result was confirmed with flow cytometry. Dashed line shows frequency distribution of alternative alleles in individuals with missing or differing flow cytometry measurement. Individuals were predicted as tetraploids, hexaploids, or octoploids when the y demonstrated an alternative allele frequency pattern of A, B or C-E, respectively.

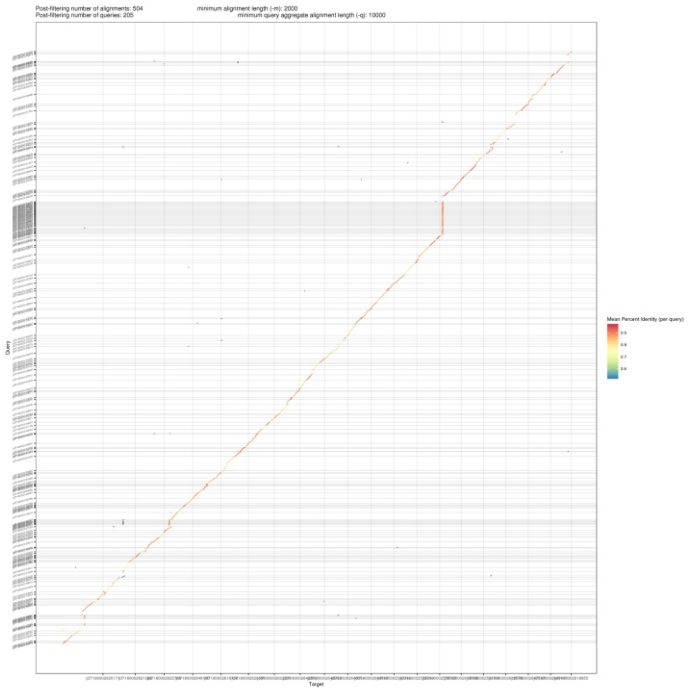


Figure S2 Syntenic alignment between Y17 and Y21, when genome of Y17 was mapped to Y21.

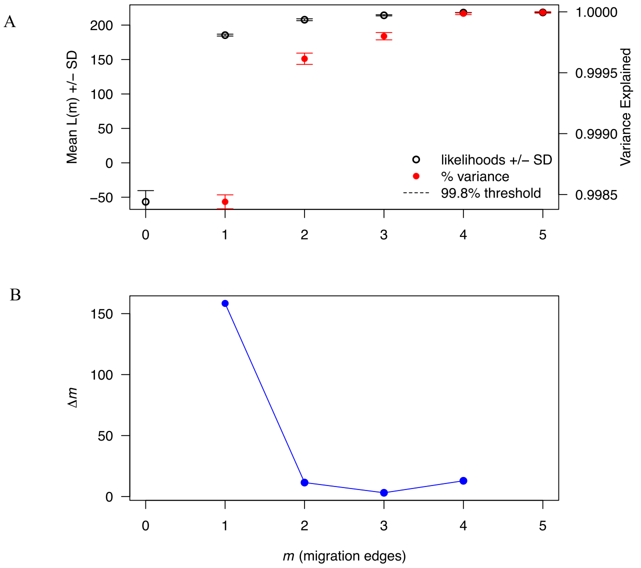


Figure S3 A)The Treemix likelihood increase trend for 1- 5 migration events, each migration event was initialized with five random initializations. The red circle and variance show the differences within each migration parameter. B) The change of maximum likelihood after adding migration edges, estimated with Evanno method.
