## Supplemental Table for "Genome-wide analysis tracks the emergence of intraspecific polyploids in *Phragmites australis*"

Table S1 Information about syntenic blocks

| | Peak 1 | | | | | Peak2 | | | | | all |
|---|---|---|---|---|---|---|---|---|---|---|---|
| | No. Blocks | Average block length | Total number of genes | Percent of genomic genes | | No. Blocks | Average block length | Total number of genes | Percent of genomic genes | | No. Blocks |
| Y17 | 914 | 23767.6 | 5154 | 3.34% | | 1063 | 19040.4 | 5637 | 3.66% | | 1977 |
| Y21 | 738 | 22518.4 | 4076 | 3.02% | | 891 | 18699.4 | 4627 | 3.43% | | 1629 |
| Y7 | 305 | 17875.4 | 1362 | 0.67% | | 311 | 14554.3 | 1363 | 0.67% | | 616 |

Table S2 Correspondance of chloroplast haplotype and lineages revealed by nuclear genomes

| Nuclear lineage | Chloroplast Haplotype | Geographic location | Reference |
|---|---|---|---|
| USnat | A-H, S, Z, AA | North America | Saltonstall., 2002 |
| AU | P,U | Eastern and Southern China, Australia | Liu et al., 2020 |
| CN | O | Northwest China | Liu et al., 2020 |
| Med | I | Gulf Coast of North America, South America, Southern Pacific, Asia | Saltonstall., 2002 |
| Usland | I ? | Gulf Coast of North America | Saltonstall., 2002 |
| EU | M | | |
| Neoploid (South Africa) | | | |


Saltonstall, K., 2002. Cryptic invasion by a non-native genotype of the common reed, Phragmites australis, into North America. Proceedings of the National Academy of Scienc

Liu, L.L., Yin, M.Q., Guo, X., Wang, J.W., Cai, Y.F., Wang, C., Yu, X.N., Du, N., Brix, H., Eller, F. and Lambertini, C., 2020. Cryptic lineages and potential introgression in


Table S3 Genetic introgression of the five ingroup lineages evaluated by ABBABABA tests

| H1 | H2 | H3 | No. Test | No. Significant(\|x\|>3) | ABBA excess(H2-H3) | BABA excess(H1-H3) | Gene flow |
|---|---|---|---|---|---|---|---|
| EU | Med | AU | 54 | 54 | 54 | 0 | Med-AU |
| EU | Usland | AU | 64 | 64 | 0 | 64 | EU-AU |
| EU | CN | AU | 64 | 64 | 64 | 0 | CN-AU |
| Med | EU | AU | 12 | 12 | 0 | 12 | EU-AU |
| Med | CN | AU | 65 | 65 | 65 | 0 | CN-AU |
| Usland | CN | AU | 64 | 64 | 64 | 0 | CN-AU |
| Med | Usland | AU | 64 | 64 | 0 | 64 | Med-AU |
| EU | Med | CN | 52 | 52 | 52 | 0 | Med-CN |
| EU | Usland | CN | 64 | 64 | 0 | 64 | EU-CN |
| Med | EU | CN | 12 | 12 | 0 | 12 | Med-CN |
| Med | Usland | CN | 64 | 64 | 0 | 64 | Med-CN |
| Med | Usland | EU | 64 | 64 | 0 | 64 | Med-EU |
| EU | Usland | Med | 64 | 57 | 0 | 57 | EU-Med |
| EU | Med | Usland | 52 | 52 | 52 | 0 | Med-Usland |
| Med | EU | Usland | 12 | 12 | 0 | 12 | Med-Usland |
| No. Med-AU | 118 | | | | | | |
| No.EU-AU | 76 | | | | | | |
| No. CN-AU | 193 | | | | | | |
| No. Med-CN | 128 | | | | | | |
| No. EU-CN | 64 | | | | | | |
| No. Med-EU | 121 | | | | | | |
| No. Med-Usland | 64 | | | | | | |
